# Functional organization of frontoparietal cortex in the marmoset investigated with awake resting-state fMRI

**DOI:** 10.1101/2021.05.31.446481

**Authors:** Yuki Hori, Justine C. Cléry, David J. Schaeffer, Ravi S. Menon, Stefan Everling

## Abstract

Frontoparietal networks contribute to complex cognitive functions in humans and macaques such as working memory, attention, task-switching, response suppression, grasping, reaching, and eye movement control. However, little is known about the organization of frontoparietal networks in the New World common marmoset monkey (*Callithrix jacchus*) which is now widely recognized as a powerful nonhuman primate experimental animal. In this study, we employed hierarchical clustering of interareal BOLD signals to investigate the hypothesis that the organization of the frontoparietal cortex in the marmoset follows the organizational principles of the macaque frontoparietal system. We found that the posterior part of the lateral frontal cortex (premotor regions) was functionally connected to the anterior parietal areas while more anterior frontal regions (frontal eye field (FEF)) were connected to more posterior parietal areas (the area around lateral intraparietal area (LIP)). These overarching patterns of inter-areal organization are consistent with a recent macaque study. These findings demonstrate parallel frontoparietal processing streams in marmosets and support the functional homologies of FEF-LIP and premotor-anterior parietal pathways between marmoset and macaque.

**Significant Statement:** Frontoparietal networks contribute to many cognitive functions in humans and macaques, but little is known about the organization of frontoparietal networks in the New World common marmoset monkey. Here, we investigated the hypothesis that the organization of the frontoparietal cortex in the marmoset follows the organizational principles of the macaque frontoparietal system. These frontoparietal connection showed overarching pattern of interareal organization consistent with Old world monkeys. These findings demonstrate parallel frontoparietal processing streams in marmosets.

## Introduction

The frontoparietal cortex in Old World primates is involved in a plethora of cognitive functions, including working memory, attention, task-switching, response suppression, grasping, reaching, and eye movement control (Andersen 1989; Andersen and Cui 2009; Barash et al. 1991; Bisley and Goldberg,M.E. 2003; Bonini et al. 2012; Caminiti et al. 1996; Colby et al. 1996; Gharbawie et al. 2011; Marconi 2001; Munoz and Everling 2004). The organization of these networks has been intensively studied in macaque monkeys using anatomical tracers, electrophysiological recordings, lesioning, electrical microstimulation, and functional magnetic resonance imaging (fMRI). Anatomical studies have shown that the frontoparietal cortex contains multiple parallel processing streams (Goldman-Rakic 1988; Petrides and Pandya 2006). Different processing models have emphasized either the segregation of these streams (Caminiti et al. 2015) or the convergence of parietal connections in certain parts of frontal cortex (Wise et al. 1997). Other models have emphasized a dorsal-ventral distinction in the frontal-parietal organization (Hoshi and Tanji 2007; Pandya et al. 2015). In addition, a “core-shell organization” has been reported in which progressively more posterior parietal regions connect to progressively more anterior frontal regions (Caspers et al. 2011).

While frontoparietal networks have been extensively studied in macaque monkeys, little is known about their organization in the New World common marmoset monkey *(Callithrix jacchus)* which is now widely recognized as a powerful nonhuman primate experimental animal that may be able to bridge the gap between humans and preclinical rodent models. The marmoset may be an ideal primate model for studying frontoparietal networks, because – unlike in the macaque monkey – the marmoset’s lissencephalic (smooth) cortex allows array and laminar electrophysiological recordings (Feizpour et al. 2021; Ghahremani et al. 2019; Johnston et al. 2019; Ma et al. 2020; Selvanayagam et al. 2019) and calcium imaging (Ebina et al. 2018; Kondo et al. 2018; Wakabayashi et al. 2018; Yamada et al. 2016) in all dorsal frontoparietal areas.

Recently, Vijayakumar and colleagues used a resting-state fMRI (RS-fMRI) connectivity and data-driven hierarchical clustering method to study the organizational principles of the macaque frontoparietal system (Vijayakumar et al. 2019). RS-fMRI is task independent, it thus does not require task matching across species and extensive training. The authors found evidence for multiple overlapping principles of organization, including a dissociation between dorsomedial and dorsolateral pathways and separate parietal–premotor and parietal–frontal pathways, demonstrating the suitability of this non-invasive approach for understanding the functional organization of the frontoparietal cortex.

Here we employed a similar approach to fully awake marmosets RS-fMRI data to investigate the hypothesis that the organization of the frontoparietal cortex in the marmoset follows the organizational principles of the macaque frontoparietal system. The results provide the foundation for electrophysiological and optical imaging explorations of frontoparietal networks in the common marmoset.

## 2 Methods

### 2.1 Animal preparation

All surgical and experimental procedures were in accordance with the Canadian Council of Animal Care policy and a protocol approved by the Animal Care Committee of the University of Western Ontario Council on Animal Care. All animal experiments complied with the Animal Research: Reporting *In Vivo* Experiments (ARRIVE) guidelines. Five common marmosets *(Callithrix jacchus,* one female; 323 ± 61 g; 1.6 ± 0.3 years old at the beginning of MRI scans) were used in this study. All marmosets underwent a surgery to implant a head chamber to fix the head during awake MRI acquisition as described in previous reports (Johnston et al. 2018; Schaeffer et al. 2019a). Briefly, the marmoset was placed in a stereotactic frame (Narishige Model SR-6C-HT), and several coats of adhesive resin (All-bond Universal Bisco, Schaumburg, Illinois, USA) were applied using a microbrush, air dried, and cured with an ultraviolet dental curing light. Then, a dental cement (C & B Cement, Bisco, Schaumburg, Illinois, USA) was applied to the skull and to the bottom of the chamber, which was then lowered onto the skull via a stereotactic manipulator to ensure correct location and orientation. The chamber was 3D printed at 0.25 mm resolution using stereolithography and a clear photopolymer resin (Clear-Resin V4; Form 2, Formlabs, Somerville, Massachusetts, USA). The marmosets were first acclimatized to the animal holder, head fixation system, and to a mock MRI environment for ~3 weeks prior to the first imaging session (Schaeffer et al. 2019a; Silva et al. 2011). Throughout the training sessions, the behavioral rating scale described by Silva et al. (2011) was used to assess the animals’ tolerance to the acclimatization procedure by the end of week 3, all five marmosets scored 1 or 2 on this assessment scale (Silva et al. 2011), showing calm and quiet behavior, with little signs of agitation.

### 2.2 MRI acquisition

Each animal was fixed to the animal holder using a neck plate and a tail plate. The animal was then head-fixed using fixation pins in the MRI room to minimize the time in which the awake animal was head fixed (Schaeffer et al. 2019a). Once fixed, a lubricating gel (MUKO SM1321N, Canadian Custom Packaging Company, Toronto, Ontario, Canada) was squeezed into the chamber and applied to the brow ridge to reduce magnetic susceptibility.

Data were acquired using a 9.4 T 31 cm horizontal bore magnet (Varian/Agilent, Yarnton, UK) and Bruker BioSpec Avance III console with the software package Paravision-6 (Bruker BioSpin Corp, Billerica, MA), a custom-built high-performance 15-cm-diameter gradient coil with 400-mT/m maximum gradient strength (Handler et al. 2020), and a 5-channel receive coil (Schaeffer et al. 2019a). Radiofrequency transmission was accomplished with a quadrature birdcage coil (12-cm inner diameter) built in-house. All imaging was performed at the Centre for Functional and Metabolic Mapping at the University of Western Ontario.

Functional images were acquired with 6-22 functional runs (600 volumes each) for each animal in the awake condition (total 53 runs), using gradient-echo based single-shot echo-planar imaging sequence with the following parameters: TR = 1500 ms, TE = 15 ms, flip angle = 40°, field of view (FOV) = 64 × 64 mm, matrix size 128 × 128, voxel size 0.5 mm isotropic, slices = 42, generalized autocalibrating parallel acquisition (GRAPPA) acceleration factor (anterior-posterior) = 2. A T2-weighted image (T2w) was also acquired for each animal using rapid imaging with refocused echoes (RARE) sequences with the following parameters: TR = 5500 ms, TE = 53 ms, FOV = 51.2 × 51.2 mm, matrix size = 384 × 384, voxel size = 0.133 × 0.133 × 0.5 mm, slice 42, bandwidth = 50 kHz, GRAPPA acceleration factor (anterior-posterior) = 2.

### 2.3 Image preprocessing

Data was preprocessed using FSL software (Smith et al. 2004). Raw MRI images were first converted to Neuroimaging Informatics Technology Initiative (NIfTI) format (Li et al. 2016). Brain masks for *in-vivo* images were created using FSL tools and the National Institutes of Health (NIH) T2w brain template (Liu et al. 2018). For each animal, the brain-skull boundary was first roughly identified from individual T2w using the brain extraction tool (BET) with the following options; radius of 25-40 and fractional intensity threshold of 0.3 (Smith 2002). Then, the NIH T2w brain template was linearly and non-linearly registered to the individual brain image using FMRIB’s linear registration tool (FLIRT) and FMRIB’s nonlinear registration tool (FNIRT) to more accurately create the brain mask. After that, the brain was extracted using the brain mask. RS-fMRI images were corrected for motion using FSL’ tool (FLIRT_ACC). Principal component analysis (PCA) was applied to remove the unstructured noise from the RS-MRI time course, followed by independent component analysis (ICA) with the decomposition number of 200 using Multivariate Exploratory Linear Optimized Decomposition into the Independent Components (MELODIC) module of the FSL software package. Obtained components were classified as signal or noise (such as eye movement, CSF pulsation, heart rate, and respiratory artifacts) based on the criteria as shown in a previous report (Griffanti et al. 2017), and noise components were regressed out from the rfMRI time course using FSL tool (fsl_regfilt). All RS-fMRI images were finally normalized to the NIH template using rfMRI-to-T2w and T2w-to-template transformation matrices obtained by FLIRT and FNIRT, followed by spatial smoothing by Gaussian kernel with the full width of half maximum value of 1.0 mm.

### 2.4 Frontoparietal functional connectivity

We first examined the relative strengths of functional connectivity (FC) of parietal areas (AIP, LIP, MIP, OPt, PE, PEC, PF, PFG, PG, PGM, and VIP) with each lateral frontal cortex (LFC) area (45, 46D, 46V, 6DC, 6DR, 6Va, 6Vb, 8C, 8aD, 8aV, 8b, and 9) and vice versa. To this end, we extracted the time courses for each run from each LFC and parietal volume-of-interest (VOI) as defined in the Paxinos marmoset atlas (Paxinos et al. 2012) registered to the NIH template (Liu et al. 2018), then a correlation map (transformed to Z-scores) was calculated with each other voxel within the brain (with mean white matter and cerebral spinal fluid time courses as nuisance regressors) using FSL’s FEAT. These z-score maps were averaged across runs for each animal, then across animals. Resultant maps were masked using LFC VOI for parietal correlations, and parietal VOI for LFC correlations, respectively, and normalized to be the maximum value=1 to show the relative connectivity strength. We also calculated the average values across animals for each VOI and created spider plots using a MATLAB toolbox developed by the Mars lab (Mr Cat; http://www.rbmars.dds.nl/lab/toolbox.html).

### 2.5 Comparison with tracer-based cellular connectivity

To directly compare the frontoparietal FC with structural connectivity, we used retrograde tracer-based cellular connectivity maps in volume space (Majka et al. 2020), which are publicly available from the marmoset brain connectivity website (http://www.marmosetbrain.org). We explicitly focused on the five regions (frontal areas 45, 6DR, 6Va, 8aD, and 8aV) that receive strong projections from the regions surrounding the intraparietal sulcus (IPS). All output volumetric data were projected to the right hemisphere. To compare the frontoparietal FC with structural connectivity, normalized z-score maps having the functional connections with each LFC (45, 6DR, 6Va, 8aD, and 8aV) were first thresholded at the value of 0.5 for visual purpose, then were mapped onto the NIH T2w atlas (Liu et al., 2018).

### 2.6 Hierarchical clustering

After all RS-fMRI images were in template space, the time courses were imported into MATLAB (The Mathworks, Natick, MA) for hierarchical clustering analysis. The time courses from each frontal or parietal VOI were extracted for each run and a partial correlation between each frontal and parietal VOIs (and vice versa) was calculated with mean white matter and cerebral spinal fluid time courses as nuisance regressors. The resultant values were transformed to z-scores, averaged across runs, then hierarchical clustering was conducted to extract discrete functional clusters based on the extrinsic functional connectivity (i.e. the strength of the connections to the parietal regions was used for LFC clustering, and vice versa). To conduct the LFC clustering, for instance, we calculated the Euclidean distance between every pair of the parietal FCs (z values) connected with LFC seed regions, then created the agglomerative hierarchical cluster tree based on the Euclidean distance using Matlab’s linkage function. Finally, these clusters were mapped onto the gray matter surface (Liu et al. 2018) using the Connectome Workbench (Marcus et al. 2011). To examine the relationships between each parietal and lateral frontal cluster, we calculated a connectivity matrix with correlation coefficients for each run. These connectivity matrices were first averaged across runs. We performed one-sample t-tests with Bonferroni correction for multiple comparisons (p < 1.0×10^-4^) to 12 connections (3 frontal and 4 parietal clusters).

## 3 Results

### 3.1 Functional and structural connectivity

We first sought to identify the strengths of functional connectivity of parietal areas with the different lateral frontal areas. As evident in Figs. 1 and 2, it seems that these frontoparietal connectivity maps can be classified into at least three groups: anterior parietal-dominant (areas 6DC, 6Va, 6Vb, A8c), posterior parietal-dominant (areas 6DR, 8aD, 8aV), and areas with weak parietal connections (areas 45, 46D, 46V, 8b, 9). Nomenclature for the brain areas are shown in Table 1. The FC map from area 8aV was broadly connected to the areas around the IPS. We next identified the strengths of functional connectivity of lateral frontal areas with each parietal area. As shown in Figs. 3 and 4, parieto-frontal connectivity maps can be classified into two groups: premotor-dominant (areas AIP, PE, PEC, PF, PFG) and caudal prefrontal (including 6DR)-dominant (areas LIP, MIP, OPt, PG, PGM, VIP) areas.

**Figure 1.**
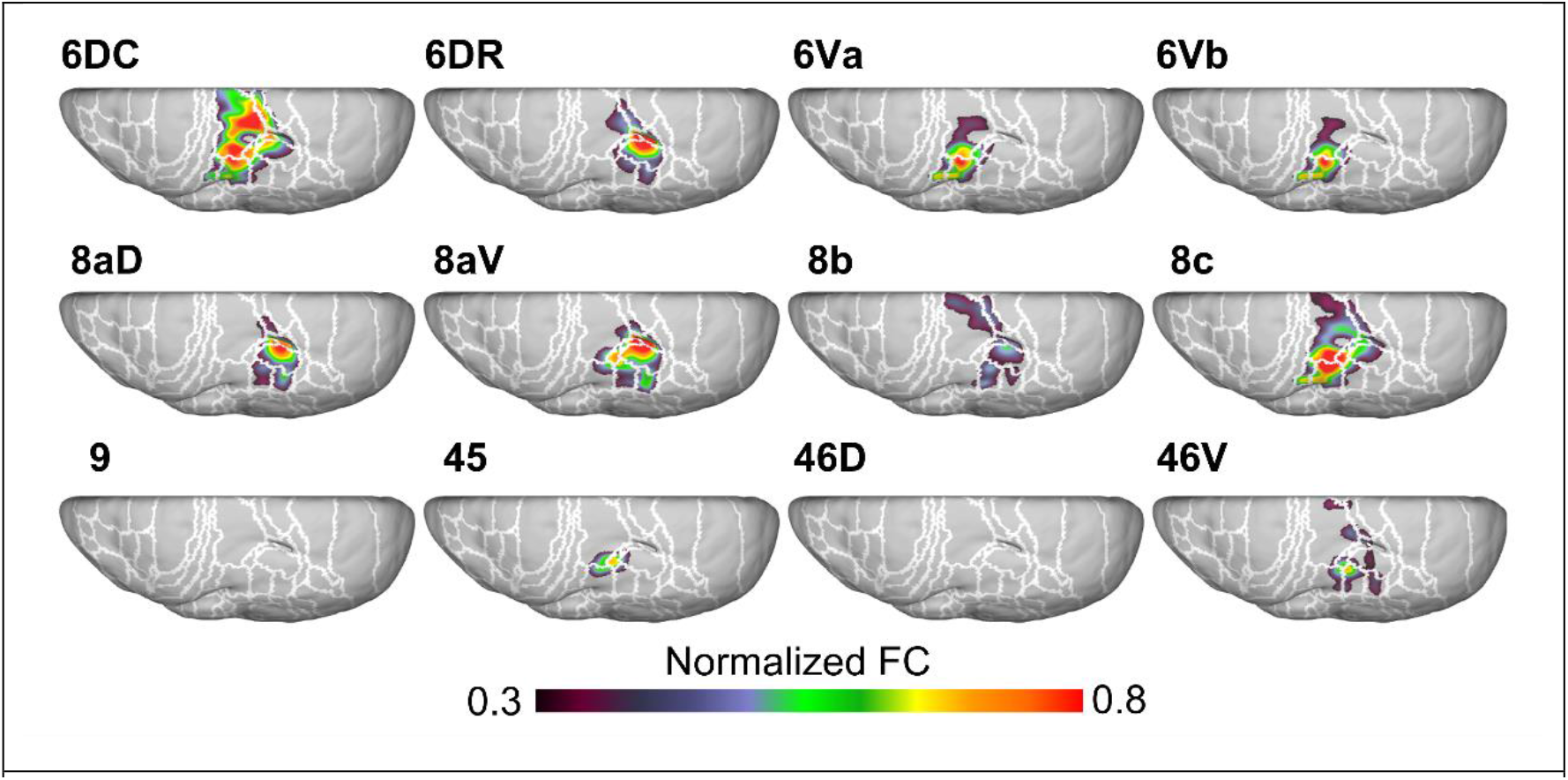
Normalized functional connectivity maps to the posterior parietal cortex from the seed of each lateral frontal area. Nomenclature for the brain areas are shown in Table 1.

**Figure 2.**
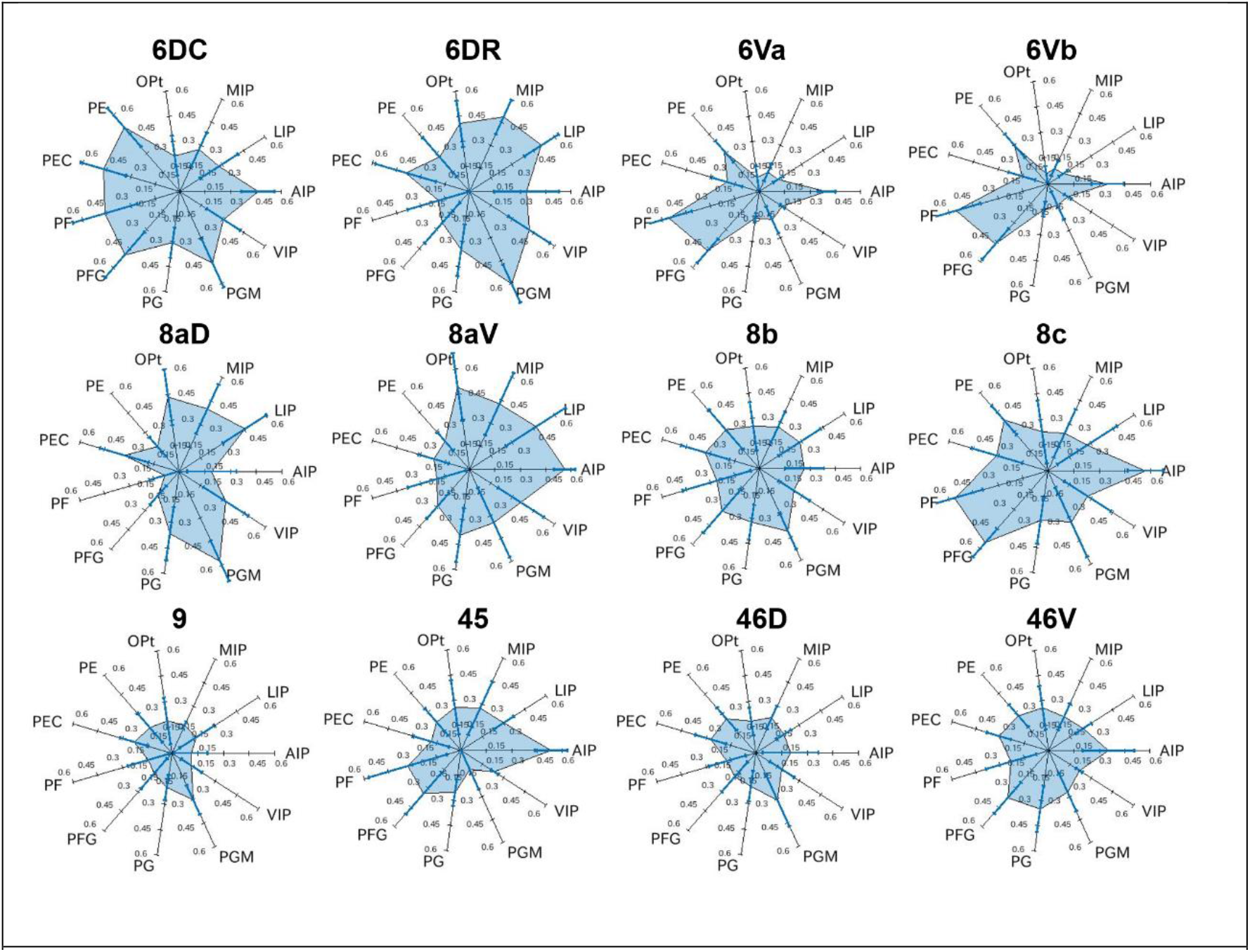
Spider plots showing parietal functional connectivity with each frontal seed region. The bars in each plot show the standard deviations among animals.

**Figure 3.**
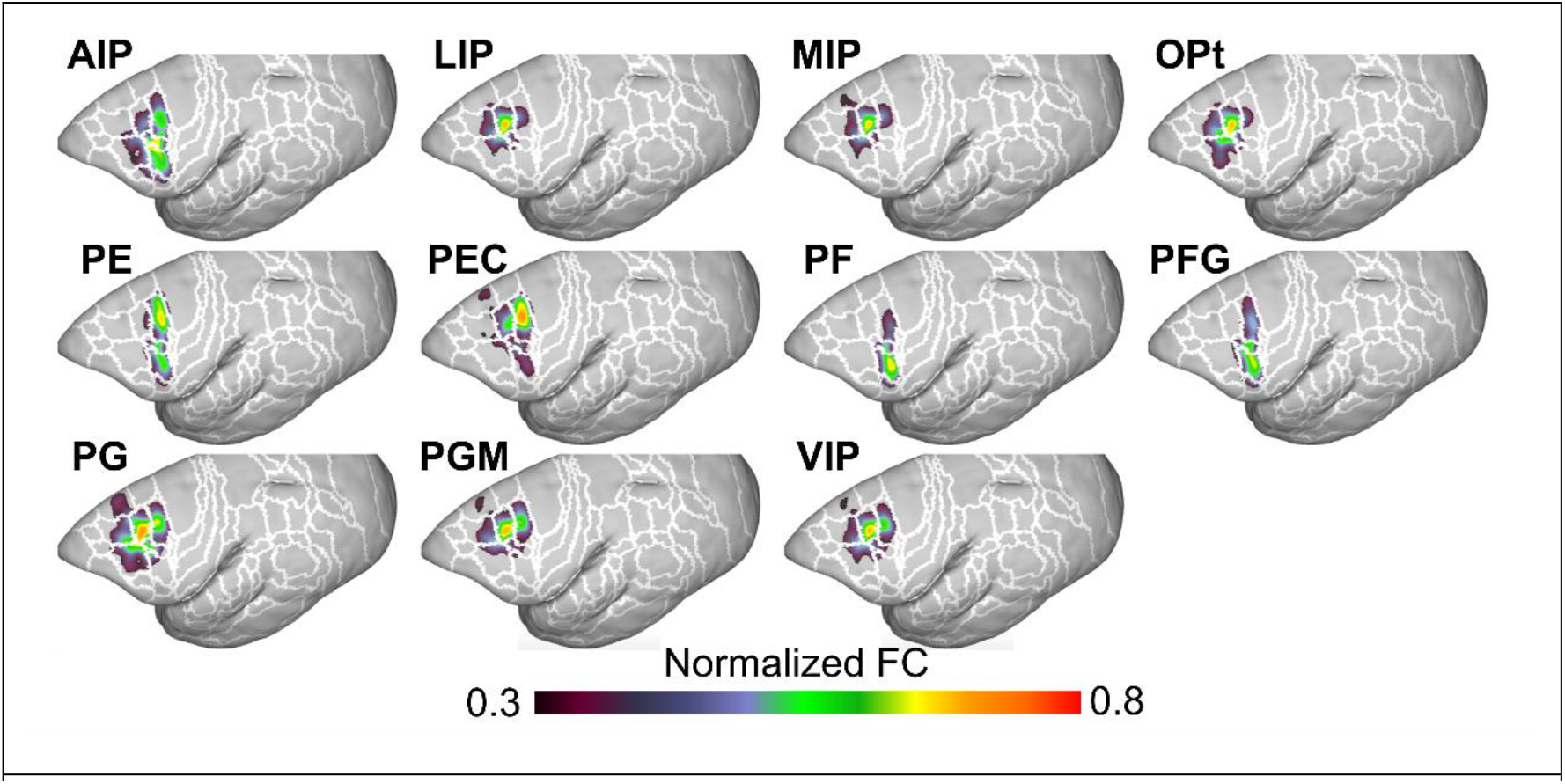
Normalized functional connectivity maps to the lateral frontal cortex (LFC) from the seed of each parietal area. Nomenclature for the brain areas are shown in Table 1.

**Figure 4.**
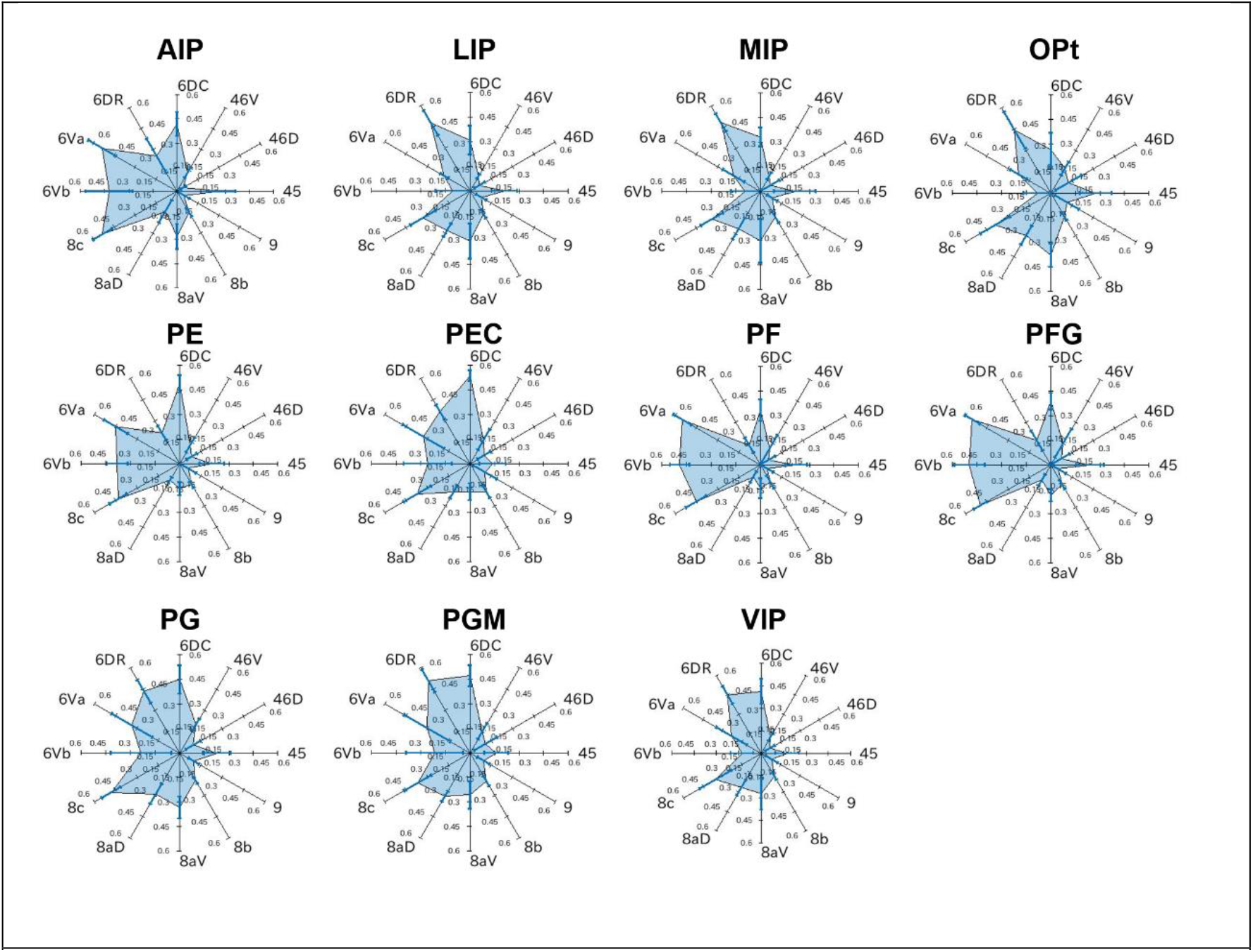
Spider plots showing lateral frontal cortex (LFC) functional connectivity with each parietal seed region. The bars in each plot show the standard deviations among animals.

**Table.**
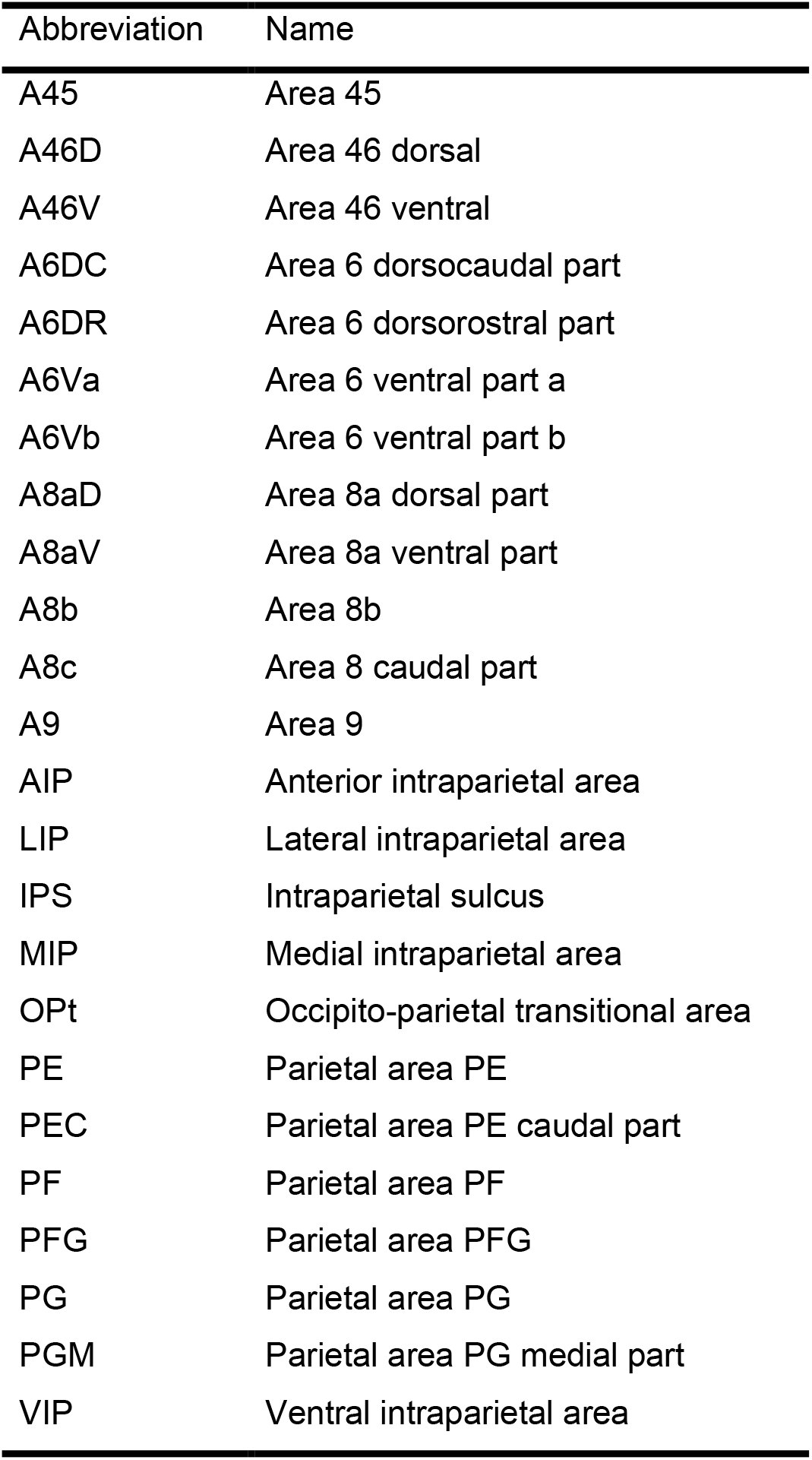

To investigate whether this tendency can be found in structural frontoparietal connections, we next compared the FC with the structural connectivity (SC) focusing on the areas around the IPS using publicly available tracer-based cellular connectivity maps (Majka et al. 2020). Areas 6Va and 45 injections showed strong structural connectivity with anterior parts of the parietal cortex, but weak connectivity with posterior parietal cortex (Fig. 5). In contrast, area 8aD and 6DR injections displayed strong structural connectivity with more posterior regions of the parietal cortex. Area 8aV was more broadly connected to a larger portion of parietal areas. These results show a good agreement between functional and structural frontoparietal connectivity in the common marmoset.

**Figure 5.**
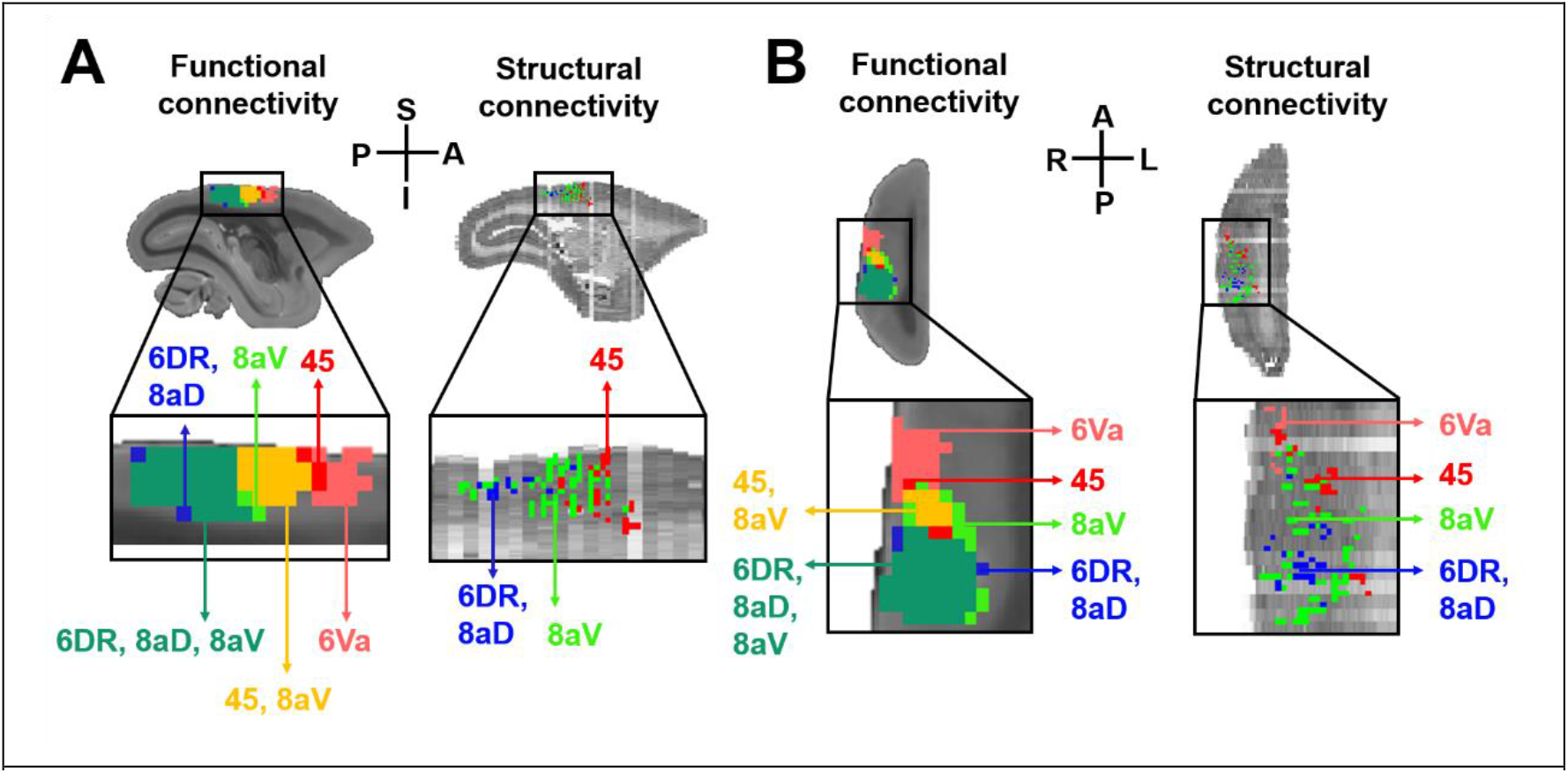
Comparison between functional and structural connections for the areas surrounding the intra parietal sulcus from each frontal region in sagittal (A) and axial views (B). Each color region in the parietal area shows the connection from 6Va (pink), 45 (red), 45 and 8aV (yellow), 8aV (light green), 6DR, 8aD and 8aV (dark green), and 6DR and 8aD (blue).

### 3.2 Frontoparietal Functional clustering

To systematically cluster the LFC and parietal cortex, hierarchical clustering was conducted based on the extrinsic functional connectivity (i.e., the strength of the connections to the parietal regions was used for LFC clustering, and vice versa). As shown in the hierarchical tree for the frontal cluster (Figs. 6A and B), the initial branching into LFC clusters dissociated premotor areas (6DC, 6Va, 6Vb, 8c) from the remaining part of the LFC, which then further separated into lateral (6DR, 8aD, 8aV) and medial clusters (45, 46D, 46V, 8b, 9). The initial branching into two parietal clusters separated the lateral part from the medial part (PGM) of the parietal area (Fig. 4B). The lateral part was then further separated into anterior (PE, PF, PFG), middle (AIP, PEC, PG, VIP), and posterior regions (LIP, MIP, OPt) (Figs. 7A and B).

**Figure 6.**
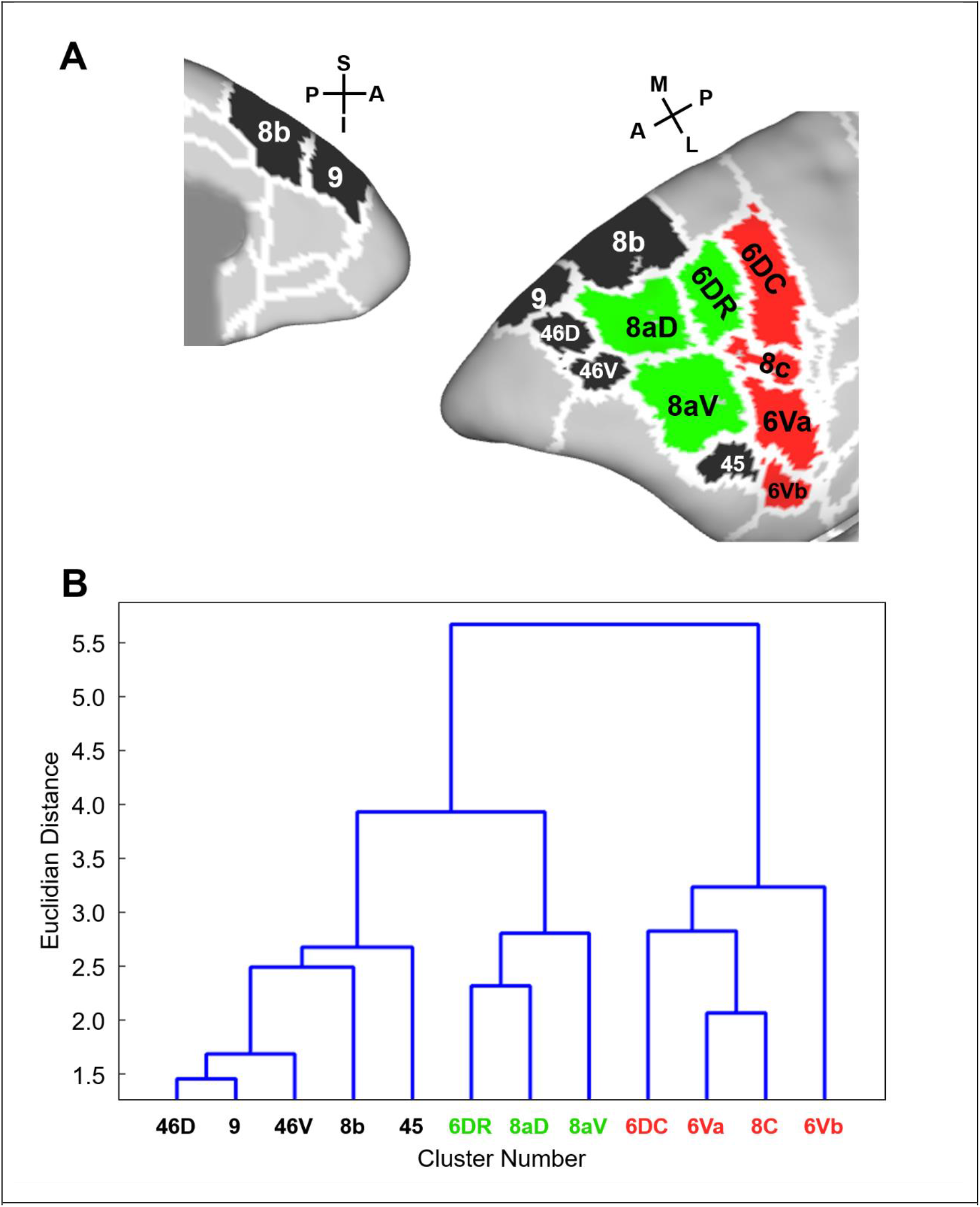
Functional clusters in the lateral frontal cortex (LFC) (A) and their dendrograms (B). The colors in each cluster (A) correspond to the colors of characters in the panel B.

**Figure 7.**
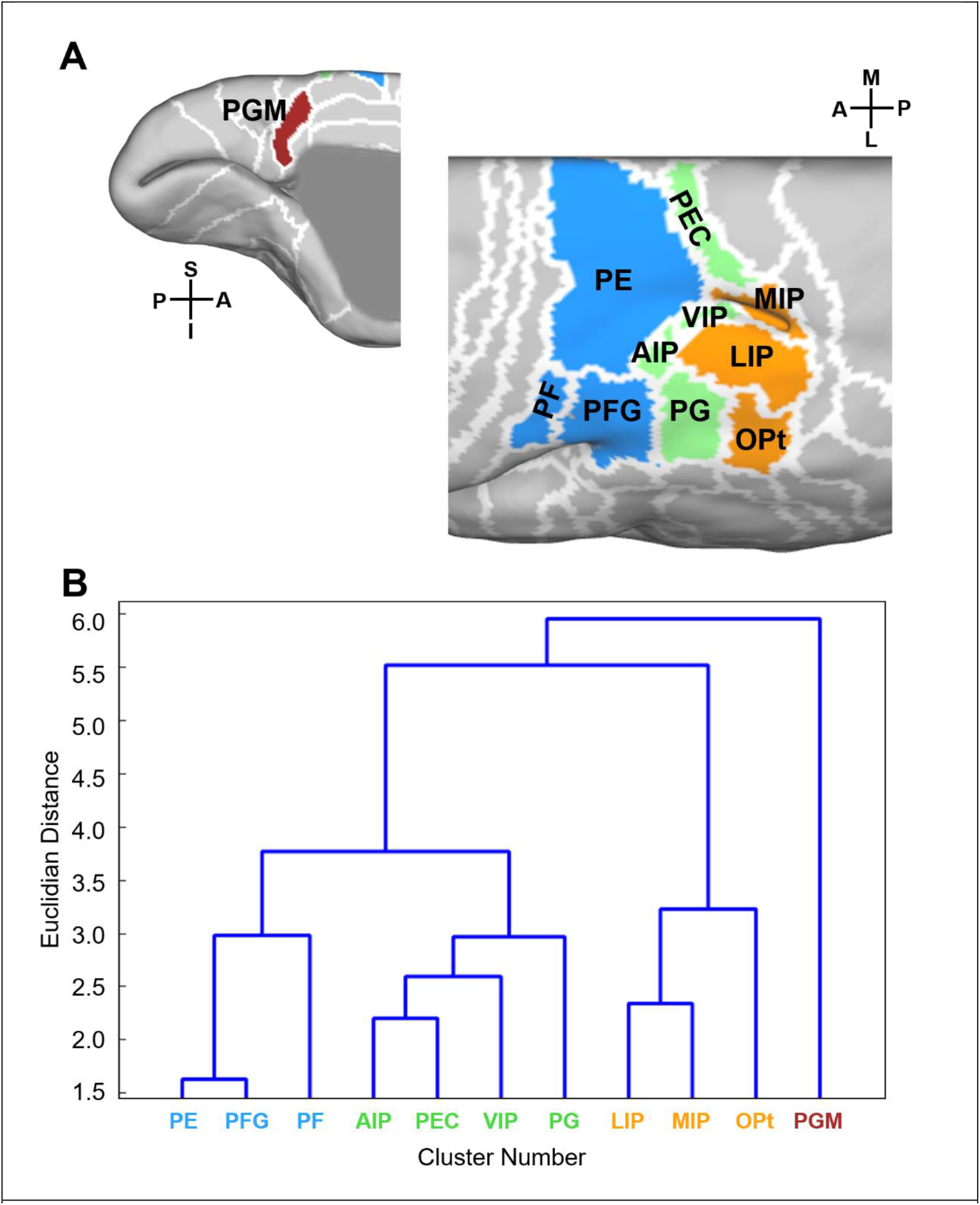
Functional clusters in the parietal regions (A) and their dendrograms (B). The colors in each cluster (A) correspond to the colors of characters in the panel B.

### 3.3 Relationship between frontal and parietal clusters

To examine frontoparietal functional connectivity, we calculated a correlation matrix that describes the average connectivity strength between frontal and parietal clusters (Fig. 8A). The strongest connections between lateral frontal and parietal clusters are plotted in Fig. 8B. These frontoparietal connections showed that the posterior part of the LFC was functionally connected to anterior parietal areas while the relatively anterior part of LFC was functionally connected to more posterior parietal areas, showing an overarching patterns of inter-areal organization. Moreover, the anterior part of LFC was also strongly functionally connected to medial parietal cortex. LPFC cluster 1 which includes area 8b, 9, 46D and 46V did not exhibit strong functionally connectivity with parietal regions.

**Figure 8.**
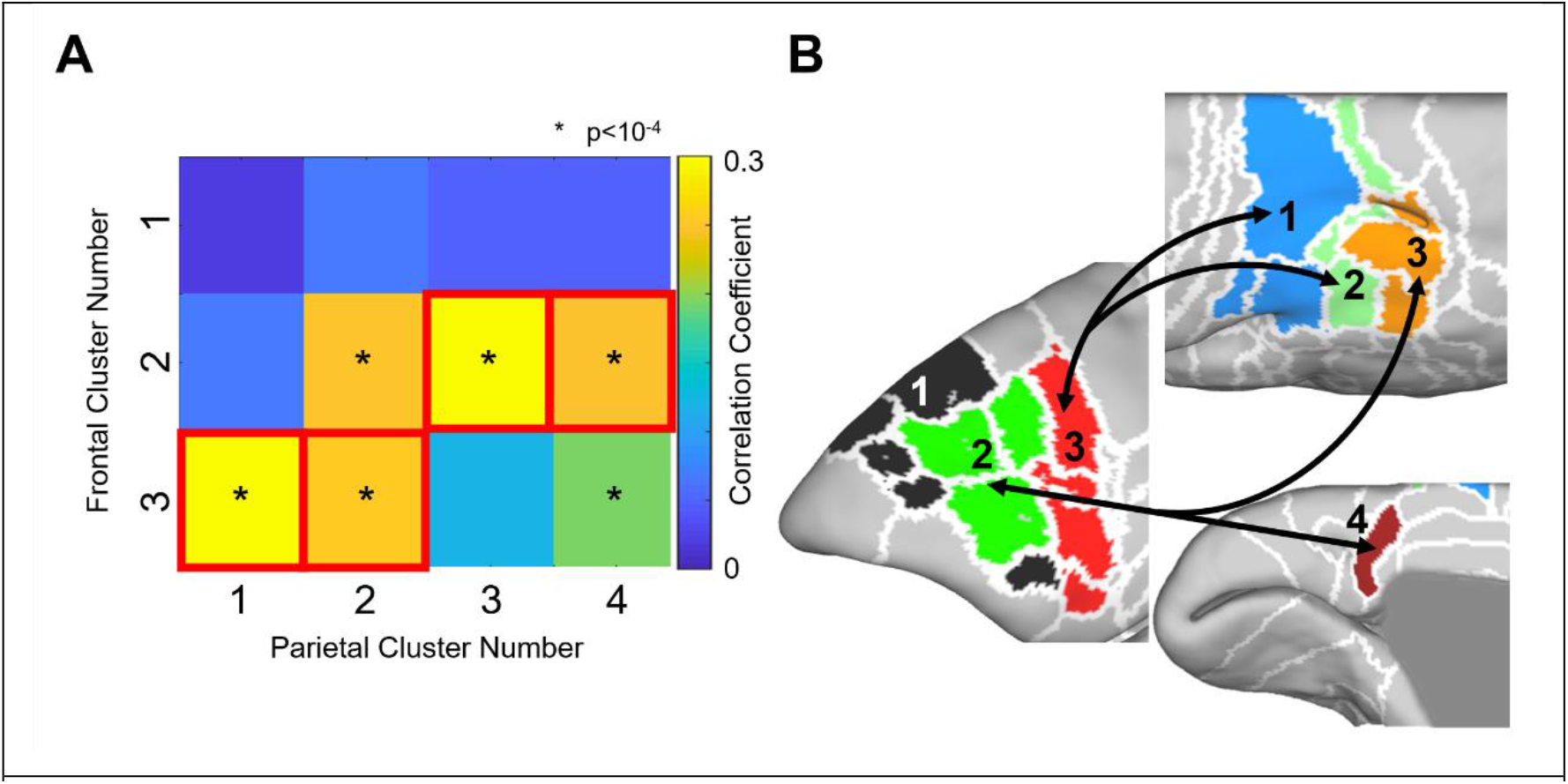
Functional connectivity matrix of the 3 frontal (rows) and 4 parietal clusters (columns), showing the average connectivity strength across two lobes. The red squares show the strongest connections to the lateral frontal area from each parietal cluster and vice versa (A). Clusters were overlaid on the marmoset surface with arrows indicating the strongest functional connections (B).

## 4 Discussion

In this study, we aimed to test the hypothesis that the organization of the frontoparietal cortex in the marmoset follows similar organizational principles as the macaque frontoparietal system. To this end, we used RS-fMRI and clustered the posterior parietal cortex (PPC) and LFC based on the strength of the functional connectivity between the regions. The initial branching into two parietal clusters separated the lateral part from the medial part (PGM) of the parietal area. The lateral part was then further separated into anterior (PE, PF, PFG), middle (AIP, PEC, PG, VIP), and posterior regions (LIP, MIP, OPt). The initial branching into LFC clusters dissociated premotor areas (6DC, 6Va, 6Vb, 8c) from the remaining part of the LFC, which then further separated into lateral (6DR, 8aD, 8aV) and medial clusters (45, 46D, 46V, 8b, 9). The anterior and middle parts of PPC was connected to premotor area, whereas the posterior and medial parts of PPC were relatively connected to caudal prefrontal areas (6DR and 8a). These overarching patterns of inter-areal organization are consistent with a recent macaque study using RS-fMRI and a hierarchical clustering technique (Vijayakumar et al. 2019). Macaque anterior parietal lobe areas (PF, PG) and AIP are functionally connected to the ventral premotor area, whereas the posterior parietal lobe (OPt) and IPS (AIP, VIP, PEa) are connected to the caudal dorsolateral cortex. In macaques, premotor and anterior parietal regions are known to be involved in the reaching and grasping systems (Bonini et al. 2012; Caminiti et al. 1996; Gharbawie et al. 2011; Marconi 2001), while dorsal prefrontal and posterior inferior parietal lobe (IPL) areas are known to be involved in the oculomotor control and spatial attention in macaques (Andersen 1989; Andersen and Cui 2009; Barash et al. 1991; Bisley and Goldberg. 2003; Colby et al. 1996; Munoz and Everling 2004).

Although less is known about the functional organization of the marmoset frontoparietal network compared to the extensively studied macaque, a few recent studies highlight functional similarities between marmosets and macaques. For example, electrical microstimulation in areas 45 and 8aV evoked small saccade eye movements, while microstimulation in 6DR and 8C evoked larger saccades with shoulder, neck and ear movements (Selvanayagam et al. 2019). These findings are consistent with the properties in ventrolateral (Bruce et al. 1985), and dorsomedial macaque FEF, respectively (Corneil et al. 2010; Elsley et al 2007). Microstimulation in marmoset area LIP (Ghahremani et al. 2019) evoked constant-vector and goal-directed saccades similar to previous macaque studies (Shibutani et al. 1984; Their and Andersen 1996, 1998). When the marmosets performed a gap saccade task, single-unit activity in the area surrounding the IPS showed a neural correlate for the gap effect (Ma et al. 2020), similar to macaque studies (Chen et al. 2013, 2016). These findings support a functional homology of FEF and LIP between marmoset and macaque. In addition, several fMRI studies also support a homologous organization of frontoparietal cortex in marmosets and macaques. Resting-state fMRI studies have identified a frontoparietal network in macaques (Hutchison et al. 2011, 2012) and marmosets (Ghahremani et al. 2016; Hori et al. 2020a) surrounding 8aV and LIP. Task-based fMRI has shown that area 8aV and a broad region of the PPC including LIP were activated during a saccade task in marmosets (Schaeffer et al. 2019b) and macaques (Koyama et al. 2004). Taken together with our findings that area 8a is more strongly functionally connected to the posterior part of PPC, these connections might play a common functional role in marmosets and macaques.

A previous human RS-fMRI study has revealed the connection maps with three main parietal regions (superior parietal lobe (SPL), anterior IPL, and posterior IPL) (Vincent et al. 2008). The SPL correlations extended from the dorsal premotor cortex (Vincent et al. 2008) and the distribution were consistent with the dorsal attention system (Thomas Yeo et al. 2011). Further anterior IPL has been shown to correlate with the anterior PFC (Vincent et al. 2008), often activated by cognitive control tasks (Botvinick et al. 2004; Dosenbach et al. 2007; Gruber and Goschke 2004; Koechlin et al. 1999; Ramnani and Owen 2004), whereas the posterior IPL was correlated with the dorsal frontal cortex, anterior and posterior cingulate cortex (Vincent et al. 2008) and showed the default mode network (Greicius et al. 2003; Raichle et al. 2001). Similar networks have also been found in marmosets, and subcortical activation in each network showed overlap between humans and marmosets (Hori et al. 2020c). Taken together with our findings, the organization of the frontoparietal network seems to be a fairly conserved feature across New (i.e. marmosets) and Old World primates (i.e. macaques and humans). However, unlike in humans and macaque monkeys, we could not find a clear superior-inferior (dorsal-ventral) distinction in the marmoset frontoparietal organization, although it has also been shown that the dorsal division of the PPC in marmosets has larger pyramidal neurons in layer 5 (Palmer and Rosa 2006) and more robust myelination compared to the ventral division (Rosa et al. 2009). Interestingly, Van Essen and Dierker (Van Essen and Dierker 2007) estimated the cortical expansion between human and macaques using anatomical MRI and 23 functional landmark constraints (Orban et al. 2004) and revealed the greatest expansion in humans in the inferior parietal lobule. Although it is still unknown whether local cortical expansion has occurred by the emergence of new areas or by differential expansion of existing areas in a common ancestor (Van Essen and Dierker, 2007), the differences of the superior-inferior (or dorsal-ventral) functional organizations between marmosets and Old World primates might be due to cortical expansion in the Old World primate lineage.

The Rosa group has extensively studied anatomical frontoparietal connections in marmosets using tracer injections. The group showed that PE and PEC are the main areas that send projections to area 6DC (Burman et al. 2014), whereas PF and PFG are the main parietal sources of projections to area 6Va (Bakola 2015; Burman et al. 2015). Although our result does not show distinct clusters between ventral and dorsal premotor pathways, areas PE and PEC showed stronger functional connectivity to dorsal than to ventral premotor areas, and PF and PFG exhibited stronger connectivity to ventral premotor cortex (Fig. 4). The Rosa group also showed the details of the afferent connections of subdivisions of frontal area 8 – the dorsal part 8aD receives relatively strong inputs from area LIP, and the ventral part 8aV receives widespread projections from the PPC (Reser et al. 2013). In contrast area 8b only receives weak projections from the PPC (Buckner and Margulies 2019; Reser et al. 2013). These anatomical studies are in good agreement with our results of functional connections, although there is no directional information in functional connectivity measures. Taken together with our previous finding that functional connectivity reflects the strength of monosynaptic pathways (Hori et al. 2020b), frontoparietal organization based on functional connectivity seems to be strongly linked to its anatomical connections. While invasive tracing techniques and ex-vivo cytoarchitecture studies are the gold standard for understanding the organization of the marmoset brain (Abe et al. 2018; Burman et al. 2006, 2014, 2015; Lin et al. 2019; Majka et al. 2016, 2021; Reser et al. 2013, 2017; Rosa et al. 2009), functional connectivity measures based on resting-state fMRI allow rapid large-scale comparative mapping of the brain organization across species including humans (Vijayakumar et al. 2019).

In summary, we used awake resting-state fMRI to identify frontoparietal functional clusters in marmosets. Like humans and macaque monkeys, the results demonstrate a coreshell frontoparietal organization. These patterns were also found in the structural frontoparietal connections, all of which support the view that this organization is largely conserved across primates. Unlike in humans and macaques, on the other hand, we could not identify superior and inferior frontoparietal subdivisions in marmosets. The marmoset’s small and smooth brain is ideal for laminar electrophysiological recordings in frontoparietal regions (Johnston et al. 2019). Our results provide the foundation for future explorations of frontoparietal networks in the common marmoset.

## Author contribution

Y.H., D.J.S., and S.E. designed research; Y.H., J.C.C., and D.J.S. performed research; Y.H. analyzed data; Y.H. wrote the paper; and Y.H., J.C.C., D.J.S., R.S.M. and S.E. edited the paper.

## Conflict of interest

The authors declare no conflict of interest.

## Notes

### Competing Interest Statement

The authors have declared no competing interest.

